# Functional organization of mouse auditory cortex in response to stimulus complexity and brain state

**DOI:** 10.1101/2022.08.11.503675

**Authors:** Navvab Afrashteh, Zahra Jafari, Jianjun Sun, Michael Kyweriga, Majid H. Mohajerani

## Abstract

The functional organization of sensory cortices is modulated by both extrinsic events and intrinsic states. The present study aimed to assess the mouse auditory cortex (AC) responses under varying conditions of stimulus complexity and brain state. Using wide-field calcium imaging, our results suggest a complete outline of topographic maps of frequency and FM rate as well as highly responsive regions to mouse ultrasonic vocalizations (USVs) in both awake and anesthetized states. Three new regions responsive to high-frequency tones and four new gradients responsive to frequency modulations (FMs) were identified. These maps are highly replicable across weeks and between animals. In awake versus anesthetized states, cortical responsiveness to pure tones was stronger, and regions that preferentially responded to slow rate FMs were smaller. In both states, fast FM regions showed the greatest contribution to the processing of USVs. Finally, our modeling of how best tone frequency or FM rate changes as a function of distance along a topographic gradient resulted in a sigmoid function. Together, our findings provide a better understanding of mouse AC functional organization and how this organization is modulated by changes in stimulus complexity and brain state. The function of newly identified regions in higher-order auditory/vocal processing and animal behavior should be considered in future research.

## 1. Introduction

Topographic maps are a prominent characteristic of the cortical surface. Throughout the nervous system structures that make up the auditory pathways, from the cochlea to the auditory cortex (AC), neurons are topographically ordered by their response to different frequencies. This tonotopic organization is characterized by a spectrum of gradients extended from neurons that preferentially respond to certain frequencies, ranging from high to low (Humphries et al., 2010). Some researchers have proposed that topographic maps play no or an insignificant role in cortical processing; that they are not necessary for cortical functions (Gu et al., 2018) or are merely an economic solution to intracortical connectivity (Chklovskii and Koulakov, 2004). Contrary to these ideas, other research supports the major contribution of topographic maps to sensory processing (Kaas, 1997), information integration (Wandell et al., 2007), and perceptual behaviors (Brewer and Barton, 2016).

The mouse AC is generally categorized into two major regions: the core and belt auditory areas (Joachimsthaler et al., 2014). The “core auditory area”, located at the center of the AC, consists of the primary auditory cortex (AI) and the anterior auditory field (AAF), with best frequency spatial gradients (tonotopic axes) that mirror each other (Hackett et al., 2011; Guo et al., 2012; Joachimsthaler et al., 2014; Tsukano et al., 2015; Tsukano et al., 2016). The core auditory area is known to receive globally tonotopic projections from the ventral part of the medial geniculate body (MGBv) of the thalamus (Winer et al., 2005; Hackett, 2011; Vasquez-Lopez et al., 2017). The “belt auditory region”, which is also known as the higher-order auditory area or secondary AC, is less-characterized in the literature and includes the secondary auditory field (AII), the ultrasonic field (UF), and the dorsoposterior field (DP). The belt auditory area surrounds the core region (Tsukano et al., 2015) and receives projections from primary and non-primary parts of MGB as well as other cortical areas (Winer et al., 2005; Hackett, 2011; Ohga et al., 2018). Whereas the core auditory area provides the first level of cortical processing of sound features, such as spatiotemporal characteristics, in a context- and attention-dependent manner (King and Nelken, 2009; Bizley and Cohen, 2013), the belt auditory region contributes to more complex auditory processing, such as higher-order novelty detection and sound-object association (Geissler and Ehret, 2004; Joachimsthaler et al., 2014), task-relevant processing (Atiani et al., 2014; Elgueda et al., 2019), and decision-making (Tsunada et al., 2016).

It has been shown that the frequency tonotopy of the core area involves frequencies up to 45 kHz, and neurons with specific frequencies higher than 50 kHz are localized in the belt high-frequency regions (Stiebler et al., 1997). However, during the past decades, technological advancements in neuroimaging and recording have led to significant updates in the tonotopic organization of the belt area. For instance, electrophysiological measurements suggest the “UF” area should be considered as subparts of the AAF and AI (Guo et al., 2012). Using multiscale optical Ca^2+^ imaging, it was shown that the AI area includes two rostral regions that process high-frequency tones (Issa et al., 2014; Liu et al., 2019). A recent publication on AC tonotopy, using a combination of flavoprotein autofluorescence imaging and anatomical techniques, demonstrated that AI includes two areas that respond to high-frequency tones and that the dorsal part of the AI high-frequency area is a discrete region from the rest of the AI (Tsukano et al., 2015). Using wide-field calcium imaging in awake mice, Romero et al. (Romero et al., 2020) also introduced a new layout for AC, including a previously unidentified region called the ventral posterior auditory field (VPAF) that responds predominantly to low frequencies, with a small high-frequency region previously considered to be part of AI.

Many species, including mice, use vocalizations for communication and to support social interactions (Holy and Guo, 2005; Portfors, 2007; Portfors and Perkel, 2014). It has been argued that mice vocalize innately, and they have limited potential for learning vocalizations (Arriaga and Jarvis, 2013; Portfors and Perkel, 2014). Despite this limited capacity, in recent years, there has been growing interest to study the neural basis for speech and socio-cognitive disorders in this species (Enard et al., 2009; Scattoni et al., 2009; Fischer and Hammerschmidt, 2011; Portfors and Perkel, 2014). Mouse ultrasonic vocalizations (USVs) are often classified based on their spectro-temporal characteristics. Female and male mice utter USVs with similar spectro-temporal structures (Guo and Holy, 2007; Hammerschmidt et al., 2012). Neurons in the core and belt regions of AC respond to vocalizations, with higher stimuli selectivity in the belt region (Geissler and Ehret, 2004; Romanski and Averbeck, 2009; Carruthers et al., 2013; Carruthers et al., 2015). At the mesoscale level, a recent study determined a region in mouse AI that is more sensitive to USVs compared to pure tones (Issa et al., 2014).

Besides pure tones, frequency modulations (FMs) are prominent components of USVs and incorporate several spectro-temporal shapes, including upward and downward FM sweeps, and chevrons (Grimsley et al., 2011). It has been speculated that the same underlying mechanism in the cortex is shared for the processing of USVs and FMs (Carruthers et al., 2013). The abundance of FM sweeps in mouse USVs suggests the importance of FMs in auditory processing, and extensive studies have been devoted to characterizing how properties of FMs, including rate and direction, are represented in the brain. Detection of sweep direction is crucial in the processing of USVs, and despite findings that show neurons in AI selectively respond to this property of FMs (Zhang et al., 2003; Aponte et al., 2021), no evidence for differences between upward and downward FM sweep rate maps in mouse AC at the mesoscale level is available (Issa et al., 2017). The FM sweeps in mouse USVs include a wide range of rates and frequencies, and several topographic gradients have been identified that contribute to the processing of FM sweeps (Trujillo et al., 2011; Issa et al., 2017). But the complete cortical topographic map of FM sweep rate is unknown. The relationship between regions responsive to USVs and the topographic maps also has not been specified yet.

In addition, anesthesia affects brain functions at different levels, including cortical spatiotemporal dynamics at the mesoscale level (Afrashteh et al., 2021). Except for a few basic studies (Gaese and Ostwald, 2001; Guo et al., 2012), the literature lacks a good description of the impact of anesthesia on FM sweep rate maps and the representation of USVs in the mouse AC. Such a description would clarify understanding of how USVs, FMs, and other complex sounds are processed at the cortical level in awake and anesthetized states.

Wide-field optical imaging is a beneficial tool to explore neural dynamics at the mesoscale level (Ferezou et al., 2007; Mohajerani et al., 2010; Mohajerani et al., 2013; Issa et al., 2014). In this study, we used wide-field Ca^2+^ imaging through a large cranial window in a transgenic mouse that expresses the genetically encoded calcium indicator GCaMP6 (Nakai et al., 2001; Iadecola, 2004; Tian et al., 2009; Chen et al., 2013; Cramer et al., 2019). In this technique, the recorded fluorescence intensity directly reflects cortical neuronal activity. We aimed to characterize the topographic maps of tone frequencies and FM sweep rates, as well as to determine the impact of stimulus complexity (e.g., pure tones vs. USVs) and brain state (awake vs. anesthetized) on AC responses. We hypothesized that our study design could 1) provide a more accurate picture of mouse topographic maps; 2) reveal the impact of anesthesia on the topographic maps of best frequency and FM sweep rate, and the response of USVs relative to pure tones; 3) differentiate between upward and downward FM sweep rate maps; and 4) model the pattern of the best frequency changes in tonotopic maps, as well as the best FM rate changes in topographic maps of FM rate.

## 2. Methods

### 2.1. Animals and surgery

Twelve Thy1-GCaMP6s mice (stock 024275, strain C57BL/6J-Tg (Thy1-GCaMP6s) GP4.3Dkim/J; Jackson Laboratory) weighing ~25g were used in this study. In this mouse strain, the GCaMP6s expression level is stable in the cortical layers 2/3 and 5 excitatory neurons. The animals were cohoused in groups of two to four mice, given access to food and water ad libitum, and maintained on a 12:12-hrs light:dark cycle in a temperature-controlled room (21°C). At the age of 2 to 3 months, a chronic window implantation surgery was performed on the animals. Mice were housed individually after head-restraint implantation surgery. All the protocols and procedures were under the policies set forth by the University of Lethbridge Animal Care Committee and the Canadian Council for Animal Care.

Mice were anesthetized with 1.0-1.5% isoflurane throughout the induction and surgery procedures. A heating pad was used to preserve the body temperature at 37°C. A craniotomy was made on the right AC (3-4 mm in diameter). The lateral end of this cranial window was the squamosal bone, and the caudal end was 0.5 mm anterior to the lambdoid structure. After removing the skull parts for craniotomy, the dura was left intact and the cranial window was covered with a clear and clean glass coverslip as previously described (Mohajerani et al., 2013; Kyweriga and Mohajerani, 2016; Afrashteh et al., 2017). A stainless-steel head-plate was secured with metabond and dental cement over the mouse skull for head-fixation purposes. To have minimal contamination of light reflection from surgery borders with emitted light from GCaMP6s fluorophore throughout brain imaging, dental cement was mixed with carbon to yield a black color. After surgery, mice were housed in the recovery room for 4-7 days and received Baytril and Metacam for at least two days.

### 2.2. Animal handling and head-fixation habituation

After recovery, animal handling and head-fixation habituation procedures were carried out within 5-7 days. On the first day, animals were initially handled for 10-15min. Then the mice were placed within the imaging apparatus to explore the imaging platform for 10-15min. For the subsequent days, mice first received 10-15min of handling and then were head-fixed to a post on a metal plate that could be mounted onto the stage of the upright microscope. Afterward, the head-fixed animals were transferred to the imaging platform in the sound booth. The blue LED lights used for imaging calcium signals were kept on and the sound-booth door was closed. The duration of head-fixation was begun at 5min and increased 5min per day, with the longest duration of 25min to match the duration of an imaging session.

### 2.3. Wide-field mesoscale calcium imaging

Awake and isoflurane-anesthetized recordings were performed for 12 and 6 mice, respectively. Anesthetized recordings were performed with 1.0% isoflurane. A heating pad was used throughout the recordings to maintain the mice’s body temperature at 37°C. Wide-field optical imaging was set to assess cortical responses to sound stimuli. Each mouse was placed under the modified upright macroscope. The mouse head was oriented in a normal orientation without head rotation. The camera was angled 60-70 degrees from the horizontal head-fixation plate to be perpendicular to the auditory window. For the mesoscale calcium imaging data collection, 12-bit images were captured with 33.3ms temporal resolution (30 Hz frame rate) with a charge-coupled device (CCD) camera (1M60 Pantera, Dalsa, Waterloo, ON) and an EPIX E8 frame grabber with XCAP 3.8 imaging software (EPIX, Inc., Buffalo Grove, IL). Images were taken through a macroscope composed of a front-to-front lens combination (50mm and 135mm lenses; 4.2×4.2 mm^2^ field of view, 256×256 pixels or 16.4 μm per pixel). The cameras were focused ~0.5mm below the surface of the brain to minimize signal distortion effects caused by vasculature movements at the brain surface. A blue LED (470nm center, Luxeon Star LEDs Quadica Developments Inc., Alberta, Canada) equipped with an excitation filter (470±20nm, Semrock, New York, NY) was applied to excite the GCaMP6s fluorophore. The emitted light was filtered through an emission filter (542±27nm) before data collection at the CCD camera.

### 2.4. Auditory stimulation

Upright microscope imaging setup was placed inside a double-walled custom build sound-booth with at least 30dB sound attenuation for mouse hearing frequency range. All other setups including computers, sound processors, and data acquisition systems were placed outside the sound booth. Sound stimuli were generated using a Tucker-Davis Technologies Inc., RX6 processor (TDT, Alachua, FL, USA) and Matlab software (Mathworks, Natick, MA, USA). Sound stimuli were delivered to the left ear by a free-field speaker (TDT Inc., ES1) at ~200 kHz sampling frequency. Speaker was calibrated using a high-frequency microphone (PCB Piezotronics) and SigCalRP software (TDT Inc.). For all experiments, stimuli were played in random order with 50ms time duration and 5±1sec inter-stimulus intervals (ISI). Pure tones were generated at low (4 kHz), medium (11.3 kHz), and high (32 kHz) frequencies; and at low (40 dB), medium (60 dB), and high (80 dB) sound pressure levels (SPL). Each frequency-SPL pair was played 25 times per imaging session.

For all FM stimuli, frequency coverage was constant in the 2-64 kHz (5oct) range and the SPL was set at 60dB. Both upward and downward frequency directions and 5 sweep durations with logarithmic increments between 10-500ms (i.e., 10, 27, 71, 188, and 500 ms) were applied. The frequency sweep rate was calculated through the division of frequency coverage (5oct) by sweep duration. Each sweep rate and sweep direction was played 25 times per imaging session. For vocalization recordings, a set of 10 mouse USVs from Grimsley et al. (Grimsley et al., 2011) was used. Each of these sound stimuli represents a spectro-temporal category in mice USVs, including high-frequency sounds with various spectral shapes and time durations. They categorized mice USVs into 11 types namely: flat, one and two frequency step, chevron, downward and upward FMs, short duration, complex, harmonic, reverse chevron, and noisy. Here we excluded the noisy syllabus because they are just noisy versions of harmonic syllabus and are uttered rarely. Similar to pure tone and FM experiments, each of these USVs was presented 25 times per imaging session at 80dB SPL.

### 2.5. Preprocessing of imaging data

All preprocessing procedures were performed using MATLAB software (Mathworks, Natick, MA). The GCaMP optical signal has the highest correlation with hemodynamics between 0-0.4 Hz (Ma et al., 2016) and the effect of respiration arises between 10-13 Hz (Tort et al., 2018). To minimize respiratory and hemodynamic-related artifacts (Figure 1A-B), a 0.5-10 Hz band-pass filter was applied to each pixel time sequence. To make the images smoother, a two-dimensional spatial Gaussian filter with parameter σ = 1 pixel was applied to each frame. Then, the percentage change from baseline was determined (ΔF/F_0_) (Figure 1C). For each specific sound stimulus, the average of all trials of calcium responses was calculated and the average baseline recorded before stimulation onset was subtracted from each frame. To correct for the blurring effect of camera focusing, the average responses were deblurred using the Lucy-Richardson deconvolution method (Richardson, 1972; Lucy, 1974; Issa et al., 2014) with a point spread function specified as a Gaussian with parameter σ = 13 pixels.

**Figure 1.**
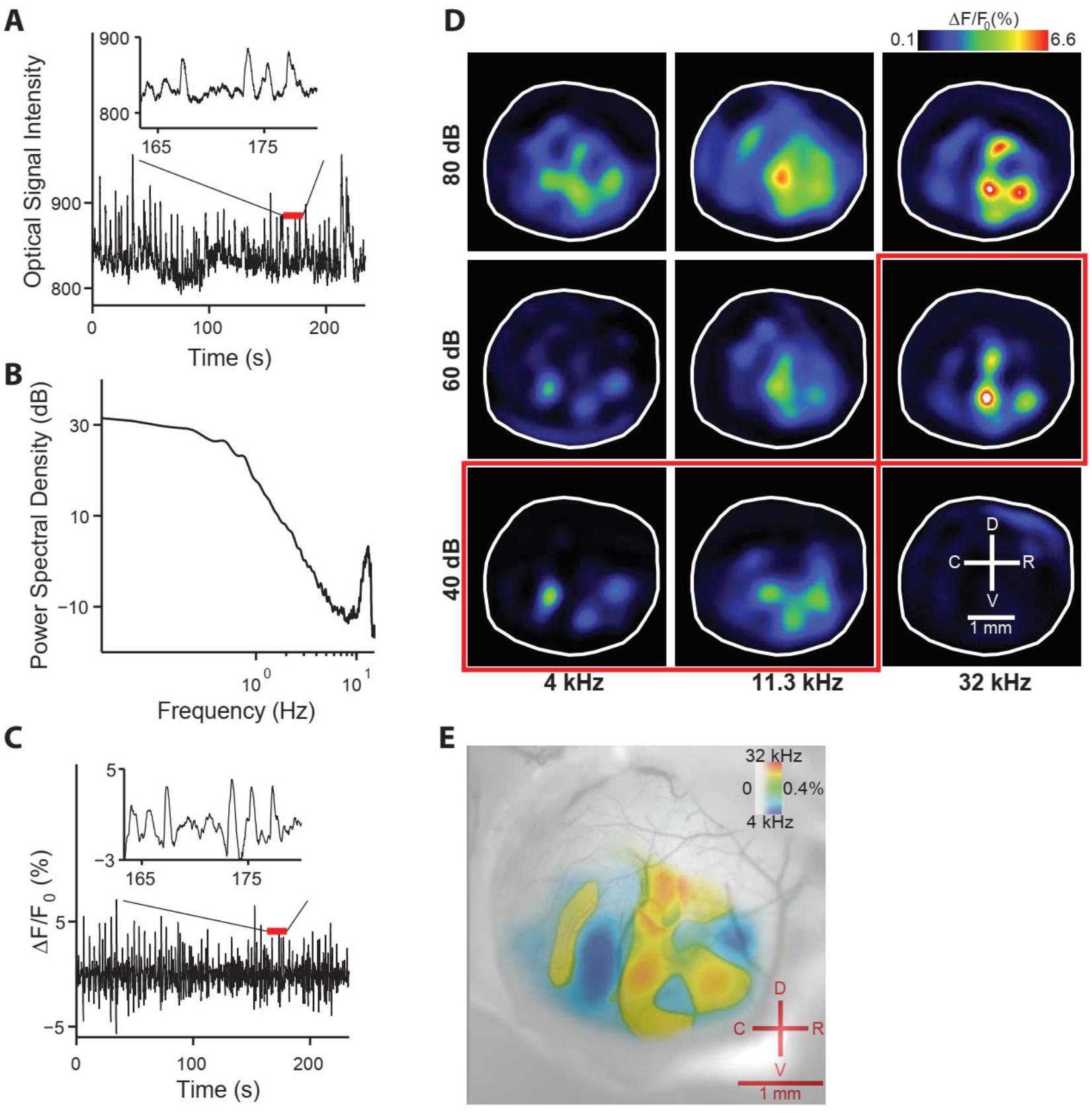
The auditory cortex responses to different tonal frequencies at three intensity levels and the resulting tonotopic map. **(A)** An example time series of raw optical signal intensity over time from a recording in the auditory cortex window. The time series consists of positive deflections that are either spontaneous or stimulus-evoked. These deflections are riding on a slow oscillation and shifted with a constant bias. The inset shows a magnified view of a shorter time window of the time series. **(B)** The power spectral density of the example optical signal intensity in **A** is plotted as a function of frequency. At ~12.75 Hz a respiration-associated oscillation results in a peak in the power spectral density. To filter out the effects of constant bias, slow oscillation, respiration, and heartbeat, the optical signal intensity is filtered through a band-pass filter from 0.5 to 10 Hz. **(C)** The filtered signal is determined in terms of the percentage change from the average baseline. The inset shows a magnified view of the time series in the same time window as in the inset plot in **A**. **(D)** The auditory cortex responses to tone presentations for an example mouse in an awake head-fixed preparation. Each of 4 kHz (low), 11.3 kHz (medium), and 32 kHz (high) frequencies were played 25 times at 40 dB (low), 60 dB (medium), and 80 dB (high) sound pressure levels. Each box exhibits the baseline subtracted average response 0-0.5 s after stimulus presentation for a frequency-amplitude pair. For each frequency, the threshold level was identified (red squares) and used to determine the tonotopic map. **(E)** The resulting tonotopic map for the example presented in section **D** is overlaid on the mouse auditory window vasculature image. The map is the average of presented frequencies weighted by responses at threshold levels and normalized by the sum of responses. The colors show the best frequency, with the saturation reflecting the magnitude of the response to the tone. Coordinates display dorsal (D), ventral (V), rostral (R), and caudal (C). The scale bar is 1 mm.

### 2.6. Tuning curves and stimuli topographic map

For pure tone experiments, tuning curves were calculated using average calcium responses for all frequency-SPL pairs (Figure 1D), and the threshold level for each frequency was quantified (red squares in Figure 1D). To generate the corresponding tonotopic maps, the top two frequencies with the highest amplitude at the threshold level in each pixel were selected. A response weighted sum of these two frequencies was determined as the best frequency at any given pixel. Pixel responsiveness also was determined proportionally to the average response amplitudes at all threshold levels. Figure 1E displays the tonotopic map calculated from the example shown in Figure 1D. In the tonotopic maps, the best frequency is shown with a color hue, in which hot colors reflect higher frequencies and cold colors represent lower frequencies. The pixel responsiveness level has been shown with color saturation, in which highly saturated colors represent pixels that are more responsive to tonal stimuli and less saturated colors show pixels with lower responsiveness to pure tones. FM sweep rate maps were determined analogously to tonotopic maps. The FM stimuli were played only at 60 dB SPL, which was used as the sound pressure level for all sweep rates.

### 2.7. Registration of topographic maps

For each animal, recording sessions across multiple days were registered based on cortical vasculatures landmarks. For tonotopic maps, low and high-frequency hotspots (11 points in total) were identified for each imaging session and similarly, for each FM experiment, slow and fast hotspots (10 points in total from upward sweep maps) were located for all FM sessions. These 21 landmarks (11 points from tonotopic maps and 10 points from FM rate maps) were used to register maps within and between animals. To register topographic maps between all animals, first, average landmarks locations for all animals were determined and used as target points in registration. The registration deformation field was constructed using landmarks from each experimental session and target landmarks as previously described in Issa et al. (Issa et al., 2014). Briefly, for each pixel, the displacement vectors of the pixel to the current session landmarks were subtracted from the displacement vectors of the pixel to the target landmarks. Then, the average difference of the displacement vectors was calculated and used to determine the estimated location of the pixel in the target reference frame.

### 2.8. Linear ridge regression analysis

To calculate the contribution of sound stimuli and other behavioral measures in the construction of the cortical signals, linear ridge regression was performed on each session of recording. The predictors include stimuli features, behavioral events, and changes in the behavioral video. Stimuli’s features for pure tones are tone frequency and sound pressure level, for FMs are sweep rate and sweep direction, and for USVs is only vocalization number. Animals’ behavior was recorded using a 3.6 mm lens Raspberry Pi infra-red (IR) camera. Five behavioral measures were used to extract behavioral events. These measures include motions in the nose, whisker pads, mouth, body, and fore paws. The behavioral events for each of the five behavioral measures were determined at times that the motion signal passed a threshold. The threshold was defined as 95 percentiles of the motion signal (equivalent to mean + 2×standard deviation but it is not sensitive to large value outliers). It was assumed that the effect of a behavioral event on the brain signals starts 0.5 s before the event onset and lasts for 2 s after the onset. To capture changes in the behavioral video that might not be included in the behavioral events, the temporal component of the first 200 principal components (PCs) of the behavioral video was also included in the regression predictors. These PCs were the outputs of applying singular value decomposition (SVD) on the vectorized video frames. The behavioral measures (motion signals) and PCs of the behavioral video were calculated using ‘facemap’ software (Stringer et al., 2019). Using SVD, the first 500 PCs of the wide-field imaging data were calculated to be used as the outputs in the linear regression modeling. To solve the linear regression, we used the code provided in Musall et al. (Musall et al., 2019) to derive the regression weights for each of the predictors.

To calculate the contribution of a set of predictors in the construction of wide-field cortical activity the explained variance (R^2^) map was determined in the following way. First, by multiplying the regression weights of the set of predictors with the values of those predictors an estimation of wide-field data is calculated. Then, squared correlation values between actual wide-field data and its estimate determine the R^2^ value at each pixel. To calculate the correlation, a window of 1 s after all stimuli onsets were used.

### 2.9. The cortical response ratio of USVs to tonal stimuli (V/T ratio)

Stimuli at 80 dB SPL were used to determine AC regions with higher responsiveness to mouse USVs compared to pure tones. First, the average responses to USVs and pure tones from all experimental sessions across multiple days were determined for each animal. Then, the average activity of each pixel was normalized to the maximum pixel value to account for GCaMP6s signal differences between animals due to possible variability in the expression level of GCaMP6s fluorophore. Responsive pixels were identified and used to exclude unresponsive pixels. A pixel was identified as responsive if its signal within a 1 s time window after stimulus onset was greater than the mean + 3×standard deviation of the baseline. After excluding unresponsive pixels, the average response to USVs was divided by the average response to pure tones (V/T) for each animal. Finally, the average V/T ratio for all animals was quantified. Note that in Figure 5B, displaying the V/T ratio, the color bar starts from 1 (V = T), and values below 1 are shown in black.

### 2.10. Statistical analyses

All statistical analyses were performed using MATLAB software. The two-sided Kruskal-Wallis test, a non-parametric alternative to analysis of variance (ANOVA), was used to evaluate if at least one variable in an array is different from others. The Chi-square goodness-of-fit test was applied to assess if values of an array come from the same distribution. The bootstrap method was also used to generate a distribution of topographic maps for each animal and session of recording. The details of bootstrapping including the number of selected trials in each iteration and the number of iterations were mentioned in the text wherever used. Statistical significance was determined at a p-value < 0.05 for all statistical analyses. To test whether the explained variance value for stimuli is greater than that of the behavioral measures and video, a one-sided Wilcoxon rank-sum test was performed. The shaded error bars used in any figure represent the standard error of the mean (SEM). For the linear model fitting of frequency or FM rate as a function of distance in a topographic gradient, the ‘fitlm’ function from MATLAB was applied and its adjusted R^2^ was reported.

## 3. Results

### 3.1. A new layout for mouse auditory cortex tonotopic map

In this experimental study, the representation of tonal stimuli over a two-dimensional surface of AC was investigated. In most previous studies on the mouse AC, the recording window was limited to a 2×2 mm^2^ area over auditory cortical regions. One of the shortcomings of using a small recording window is the likelihood of losing the response of regions close to the recording borders. In this study, a large craniotomy (3-4 mm in diameter) was created over auditory cortical regions to rule out all uncertainties associated with the size of the recording window. Three tonal stimuli representing low (4 kHz), medium (11.3 kHz), and high (32 kHz) frequencies at 40, 60, and 80 dB SPL were presented to the animal in random order. After data collection and determining the tonotopic maps for all sessions from each animal (3 awake sessions for each of the 12 animals, and one anesthetized session for each of the 6 animals), the maps were transformed into a common coordinate framework obtained from all animals’ landmark points. Figure 2A illustrates the average tonotopic maps of all awake recordings in multiple time averaging windows after the stimulus onset. Several tonotopic gradients are observed on the auditory cortical surface. The top left panel of Figure 2A exhibits a set of landmark points used in this study to register tonotopic maps onto a common framework. Some of these landmarks and their corresponding tonotopic gradients (e.g., dashed and solid lines) were previously known, and some of them were identified in this study. The previously identified tonotopic regions include AI, AII, AAF, UF (Issa et al., 2014; Tsukano et al., 2015), and the dorsal posterior region (DP) (Romero et al., 2020), and the newly characterized tonotopic regions were the dorsal anterior region (DA), the posterior auditory field (PAF), and the ventral posterior auditory field (VPAF). The VPAF region identified by Romero et al. (Romero et al., 2020) is in a different location and is primarily responsive to low frequencies. The newly identified tonotopic regions are known in the rat AC (Kalatsky et al., 2005), thus a similar terminology was used in the present study. As shown in Figure 2A, these new regions surrounded the previously identified areas and were responsive to higher frequencies. Three regions in the mouse AC, located in AI, AII, and AAF regions, were highly responsive to low frequencies. The centers of these regions are shown with the letter L in Figure 2. All three regions neighbored high-frequency regions from multiple angles, reflecting tonotopic gradients. The tonotopic gradients from the low-frequency region in AI to its high-frequency region, as well as to DP and UF have previously been described. In this study, new tonotopic gradients in AI to PAF and VPAF were recognized. These tonotopic gradients were observed dorsally or caudally to the location of the low-frequency region of AI. AI tonotopic gradients also varied in length. For instance, AI to DP was slightly greater than 1 mm, AI to PAF was approximately 0.5 mm, and AI to VPAF was slightly less than 0.5 mm. In AAF, in addition to the already known tonotopic gradients from the low-frequency region of AAF to the high-frequency region of AAF and UF, a new gradient was observed from the low-frequency region of AAF to the DA region. Similar to the tonotopic gradient into UF, this newly identified gradient was also dorsal relative to AAF, and its length (>0.5 mm) was slightly longer than the length of the main gradient in the AAF region (i.e., from the low-frequency to the high-frequency regions in AAF).

**Figure 2.**
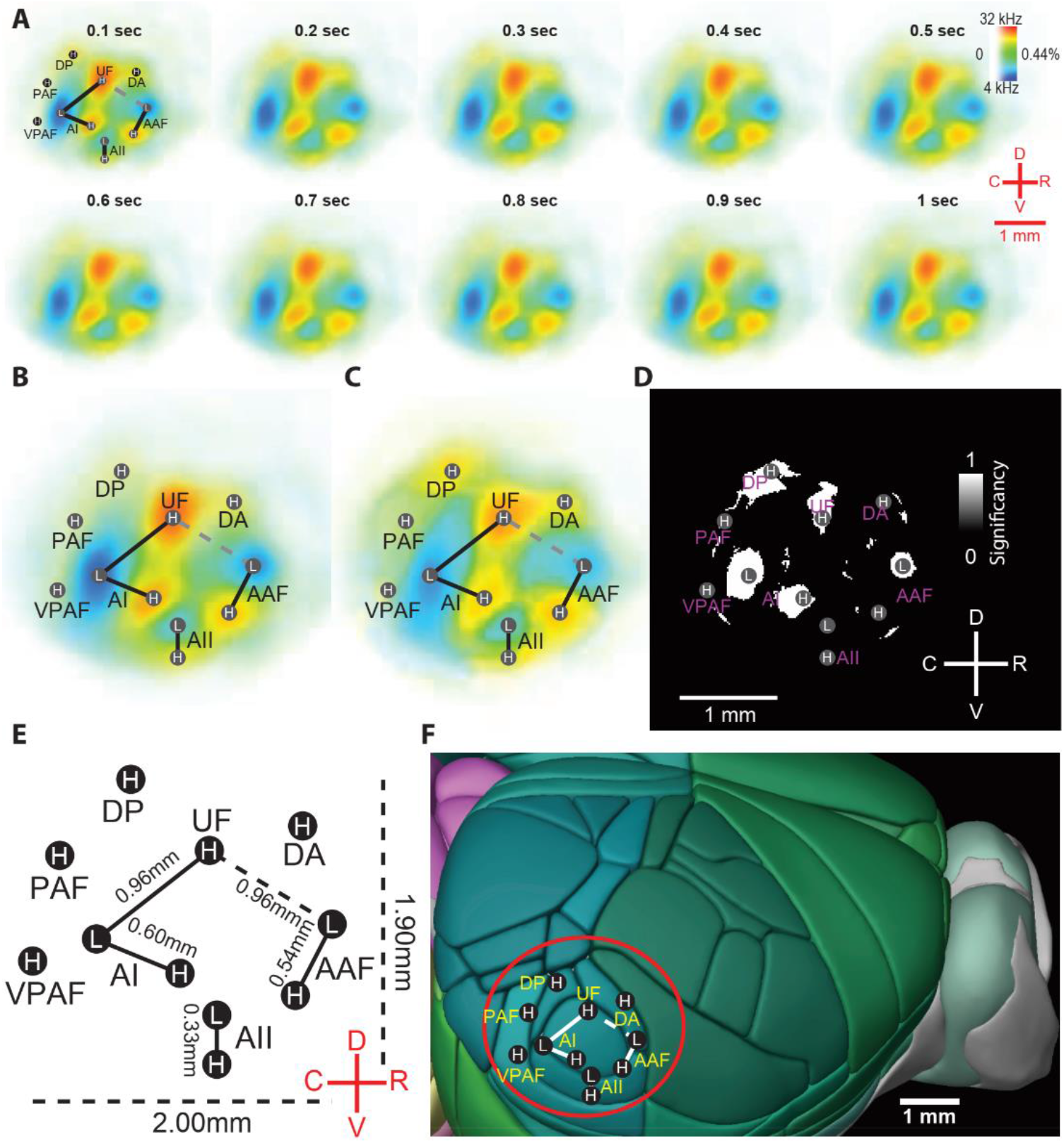
Newly identified brain areas responsive to tonal frequencies, and the impact of brain state on the tonotopic map. **(A)** The average of 36 awake and 6 anesthetized tonotopic maps from 12 three- to four-month-old Thy1-GCaMP6s mice, were merged using landmark registration. Maps of cortical responses were plotted, averaging over multiple time windows. To get the average of landmarks, maps from multiple recording sessions were aligned and merged using the transformation of landmarks (centers of low and high-frequency regions). Maps were almost identical from 100 ms after the onset of tone presentation up to 1000 ms. **(B** and **C)** The average of tonotopic maps was obtained in both awake (**B**: n = 12, recordings per mouse = 3) and isoflurane-anesthetized (**C**: n = 6, recording per mouse = 1) states. The previously identified regions (AI, AII, AAF, DP, UF) (Romero et al., 2020; Issa et al., 2014; Guo et al., 2012) and newly identified regions in this study (DA, PAF, VPAF) that respond to tonal stimuli (L for low frequencies and H for high frequencies) are presented. **(D)** Map of statistical comparison in the best frequency (the highest response to L or H frequencies) between awake and anesthetized tonotopic maps. The five white areas (AI L, AI H, AAF L, UF, and DP regions) represent regions with a significant difference (p < 0.05) in best frequency after applying the Kruskal-Wallis test at each pixel. The best frequency was significantly larger at high-frequency regions (H region of AI and UF) and reduced at low-frequency regions (L region of AI and AAF) in awake compared to anesthetized tonotopic maps. In the DP region alone, the best frequency was larger at high frequencies in anesthetized compared to awake maps. **(E)** The outline of landmarks in the mouse auditory cortex averaged from a total of 42 awake and anesthetized recordings. The auditory cortex dimensions (dorsal-ventral = 1.90 mm, rostral-caudal = 2.00 mm), and the average distance between L and H spots in AI, AII, and AAF are illustrated. **(F)** The auditory cortex tonotopic organization is overlaid on the 3D Allen Mouse Brain Atlas. Abbreviations: AI, primary auditory cortex; AAF, anterior auditory field; AII, secondary auditory field; DA, dorsal anterior field; DP, dorsal posterior field; PAF, posterior auditory field; UF, ultrasonic field; VPAF, ventral posterior auditory field.

To assess the extent of the auditory cortical regions that are more sensitive to pure tones compared to behavioral motions, a statistical test was performed between the explained variances (R^2^) of stimuli and behavioral measures and video. These explained variance maps were calculated using the outputs of a linear regression model between stimuli, behavioral measures, and behavioral video as predictors and wide-field cortical responses as outputs (see methods). The results of the statistical test of whether the R^2^ value of pure tone stimuli is larger than the R^2^ value of behavioral measures and video at each pixel are plotted in Figure S1A. Significant regions at p-value < 0.05 and the p-value map are shown on the left and right panels, respectively. At the p-value < 0.05 level, regions that show larger R^2^ for pure tones include all the high and low-frequency landmarks determined in this study. This analysis confirms that the regions identified in the current research are highly responsive to pure tones.

Figure 2A illustrates the tonotopic maps generated from multiple time averaging windows after the stimulus onset. The time duration is shown above each panel. The maps were identical starting from 100 ms after the stimulus onset. This similarity demonstrates that in the AC regions tonal stimuli evoke spatiotemporal responses that do not travel far from their initial source locations. Tone-evoked auditory cortical patterns initiated in distinct tonotopic regions then expanded radially and eventually vanished. Subsequently, the tonotopic maps with multiple time averaging windows taken from these spatiotemporal patterns were identical. Thus, tone-evoked auditory cortical responses exhibit temporally and spatially slow and stable dynamics at the mesoscale level.

Figures 2B and 2C display the tonotopic maps averaged over all animals in awake and isoflurane-anesthetized states, respectively. These tonotopic maps were calculated by averaging over a 0.5 s time window after the response onset. In both maps, low and high-frequency landmarks are represented with L and H letters, respectively. Upon visual inspection, maps in both states showed similar structures, with similar locations of hotspots, tonotopic gradients, and outlines of cortical regions. A Kruskal-Wallis test at each pixel was applied to statistically compare the two tonotopic maps in their best frequency. The statistical test results are shown in Figure 2D, in which white and black colors display regions with and without a significant difference (p-value < 0.05), respectively, between AC tonotopic maps of awake and anesthetized states. The map of corresponding p-values is plotted in Figure S1B. Five main regions showed significant differences, including the low-frequency regions of AI and AAF, and the high-frequency regions of AI, UF, and DP (median p-values shown in S1C table: AI-L p = 0.0054, AI-H p = 0.0171, AAF-L p = 0.0181, UF p = 0.0216, DP p = 0.0165). In all five regions, only a limited area located at the center showed a significant difference between awake and anesthetized states. At the center of the low-frequency regions in AI and AAF, a lower best frequency was observed in the awake map compared to the anesthetized map (note the visibly darker blue color in these regions in Figure 2B vs. 2C). At the center of the high-frequency regions of AI and UF, a higher best frequency was found in the awake versus anesthetized maps (note the darker red color at the center of both regions in Figure 2B vs. 2C). Conversely, at the center of the DP area, the anesthetized map demonstrated a higher best frequency compared to the awake map. In summary, in the awake compared to isoflurane-anesthetized condition, the sensitivity of auditory cortical regions was generally increased for both the low and high ends of the frequency spectrum (i.e., the tonotopic map expanded to lower and higher frequencies). Figure 2E exhibits the outline of the auditory cortical regions showing tonotopic organization. The distance between the centers of the two farthest regions was 2 mm on the rostral-caudal axis and 1.9 mm on the dorsal-ventral axis. Finally, Figure 2F displays the 3D atlas of the mouse brain from the Allen Mouse Brain Common Coordinate Framework (Wang et al., 2020) overlaid with the tonotopic organization of AC. The distance between the farthest landmarks in the tonotopic map supports the idea that a window larger than 2 mm in each axis is necessary to include all cortical regions involved in the AC tonotopic organization.

### 3.2. Reproducibility of tonotopic maps across time and between animals

The reproducibility of the tonotopic maps was examined in three ways: during one experimental recording session in each animal, over multiple recording sessions per animal, and between animals. To quantify the reproducibility of the tonotopic maps during a recording session in each animal, a bootstrap method was used. For 200 iterations, two random samples (out of 25 trials) were selected, and the resulting tonotopic map was calculated with the average responses of these two trials. Consequently, the distribution of tonotopic maps for the considered recording session was generated. Then, for each pixel, a Chi-square goodness-of-fit test was applied to its distribution of best frequency. Figure 3A shows the tonotopic maps calculated using the first (Figure 3A1), second (Figure 3A2), and third (Figure 3A3) parts of a recording session, with each part consisting of 8 trials (i.e., about a third of the session). Figure 3A4 represents the results of the Chi-square goodness-of-fit test on the best frequency, as explained above. In Figure 3A4, the white color indicates regions with a statistically significant (p < 0.05) change in best frequency.

**Figure 3.**
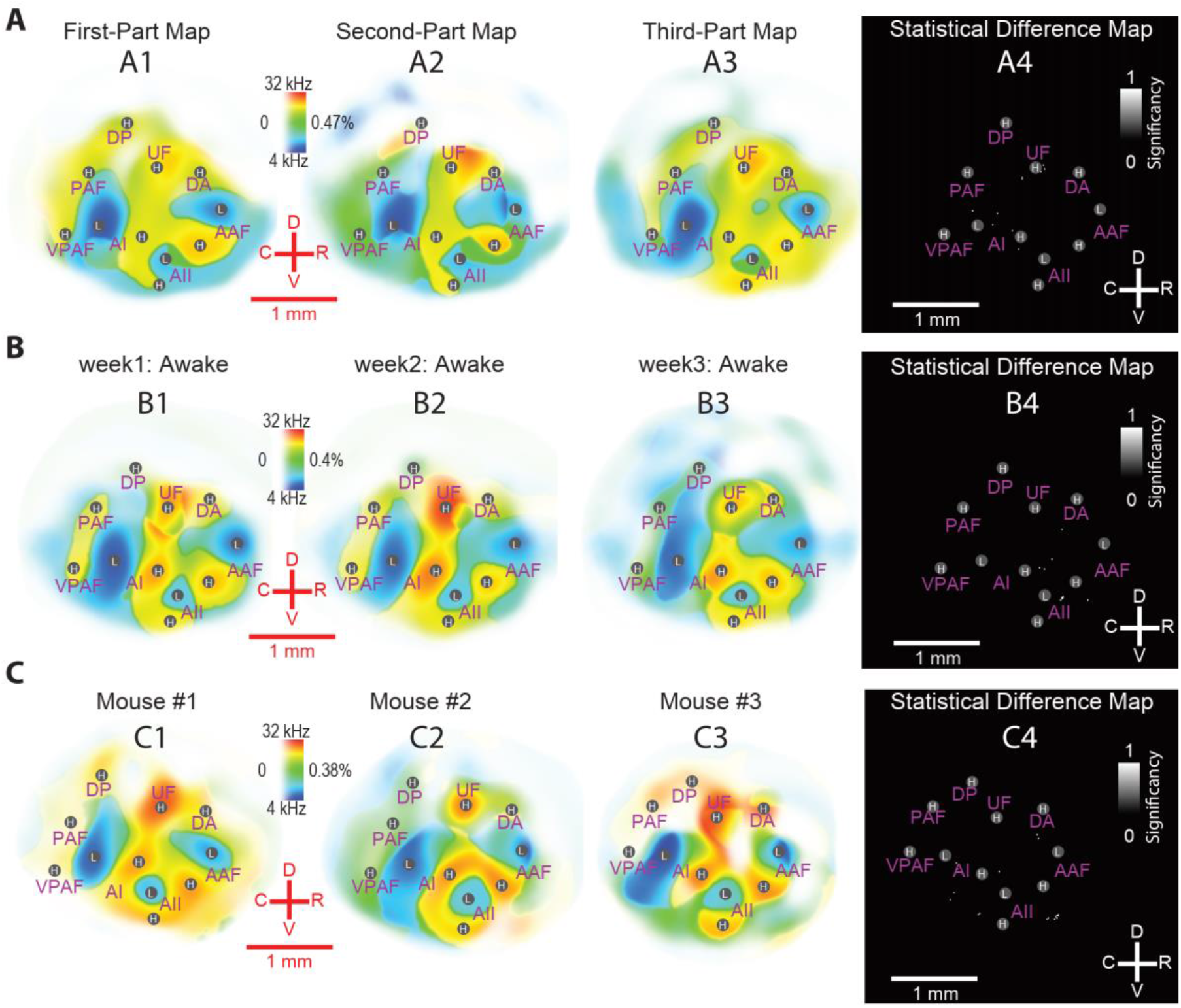
The reproducibility of mouse auditory cortex tonotopic maps was examined in three different ways including: during a recording session **(A)**, across weeks of recordings **(B)**, and between animals **(C)**. **(A)** Auditory cortex tonotopic maps derived from the first (A1), second (A2), and third (A3) parts of a recording session (8 trials per part) for an example animal in the awake state. (A4) To generate a distribution of tonotopic maps, two random trials (out of 25 trials) were bootstrapped 200 times. A Chi-squared goodness-of-fit test was applied to assess the hypothesis that the maps come from the same distribution. Except for some sparse pixels (white spots), the maps were consistent in the best frequency throughout one recording session. **(B)** Auditory cortex tonotopic maps from three consecutive weeks (B1-B3) in the awake state for the same mouse. The Kruskal-Wallis test was applied to compare the distribution of best frequencies across the three weeks after bootstrapping 20 random trials (out of 25) 100 times. B4) Except for a few sparse pixels, the tonotopic maps showed consistency over weeks of recordings. **(C)** Auditory cortex tonotopic maps from one session of recording in the awake state in 3 mice (C1-C3). The location of hotspots varied among mice, but similar low and high-frequency hot spots were shown in the 3 animals. (C4) The map of statistical analyses (the goodness-of-fit test) to examine the best frequency of all awake maps between animals (n = 12). Except for a few sparse pixels, there was no significant difference between animals in the best frequency of the tonotopic maps.

The corresponding p-value map is shown in Figure S2A. The three tonotopic maps resulting from the thirds of the experimental session were visually identical, and the goodness-of-fit test map also showed no significant differences. Overall, the structure of tonotopic gradients and the best frequency representation in the AC during an experimental session of approximately 25 min were stable and reliable.

Figure 3B shows the three resulting tonotopic maps (from all 25 trials of a session) per testing week. Visually, the three maps had a similar structure of tonotopic gradients and landmarks, with differences in the best frequency representation and/or auditory tone response sensitivity in some regions. Significant differences were assessed by generating a distribution of maps for each session using the bootstrapping method with a selection of 20 trials at each iteration for 100 iterations. Then the Kruskal-Wallis test was applied to the generated distributions of each pixel to detect if the values of the three weeks come from the same distribution. The resulting map of statistical significance (p < 0.05) is plotted in Figure 3B4 (with the corresponding p-value map in Figure S2B) and, except for a few sparse pixels, exhibits the reproducibility of the tonotopic map over weeks of recording for one animal. To examine the reproducibility of the tonotopic maps among all animals (n = 12) from the awake recordings, a goodness-of-fit test was used to test whether the best frequency at each pixel comes from the same distribution (Figure 3C). Except for a few sparse white pixels (statistical significance at p < 0.05 shown in Figure 3C4, and the corresponding p-value map in Figure S2C), the best frequency of tonotopic maps was similar across animals. Thus, AC tonotopic maps were reproducible across animals.

### 3.3. Topographic maps of FM sweep rate in awake and anesthetized states

Among a few studies of mouse AC organization in encoding FM sweep rates, the Issa et al. (Issa et al., 2017) study suffers from a small imaging window, which makes it difficult to delineate the structure of the resulting maps near the imaging borders. To evaluate the cortical responses to FM sweeps, chirp stimuli with different upward and downward sweep rates at 60 dB SPL were presented to animals in awake and isoflurane-anesthetized states. Then, FM sweep rate maps were generated for each sweep direction and brain state. Figure 4A shows the average awake and anesthetized FM sweep rate maps, examining various time windows following response onset. The maps were registered to a common framework using average landmarks of upward sweep rate maps from all animals (Figure 4A). Like the tonotopic maps, the color bar shows two properties; the color hue indicates the best FM sweep rate, and the color intensity characterizes the sound stimuli responsiveness. The extent of cortical regions responsive to FM stimuli is greater than 2×2 mm^2^, which emphasizes the importance of a large imaging window to delineate a complete topographic map. The slow and fast rate landmarks are shown with the letters S and F, respectively. The current topographic map of the FM sweep rate demonstrates a complete layout of gradients in auditory cortical regions. There are five slow and fast rate landmarks as well as sweep rate gradients distributed across auditory cortical regions. Three pairs of slow and fast rate landmarks in AI, AII, and AAF have previously been characterized (Issa et al., 2017), and two new pairs, located adjacent to the border of the imaging window, were identified in this study. All five gradients have a rostral-to-caudal direction, with the slow rate landmark more rostral than the fast rate landmark. The slow rate landmarks were in AI, AII, AAF, DA, and UF, whereas the corresponding fast rate landmarks were situated between PAF and VPAF, in AII, between AII and AAF, between AAF and UF, and between DP and PAF, respectively.

**Figure 4.**
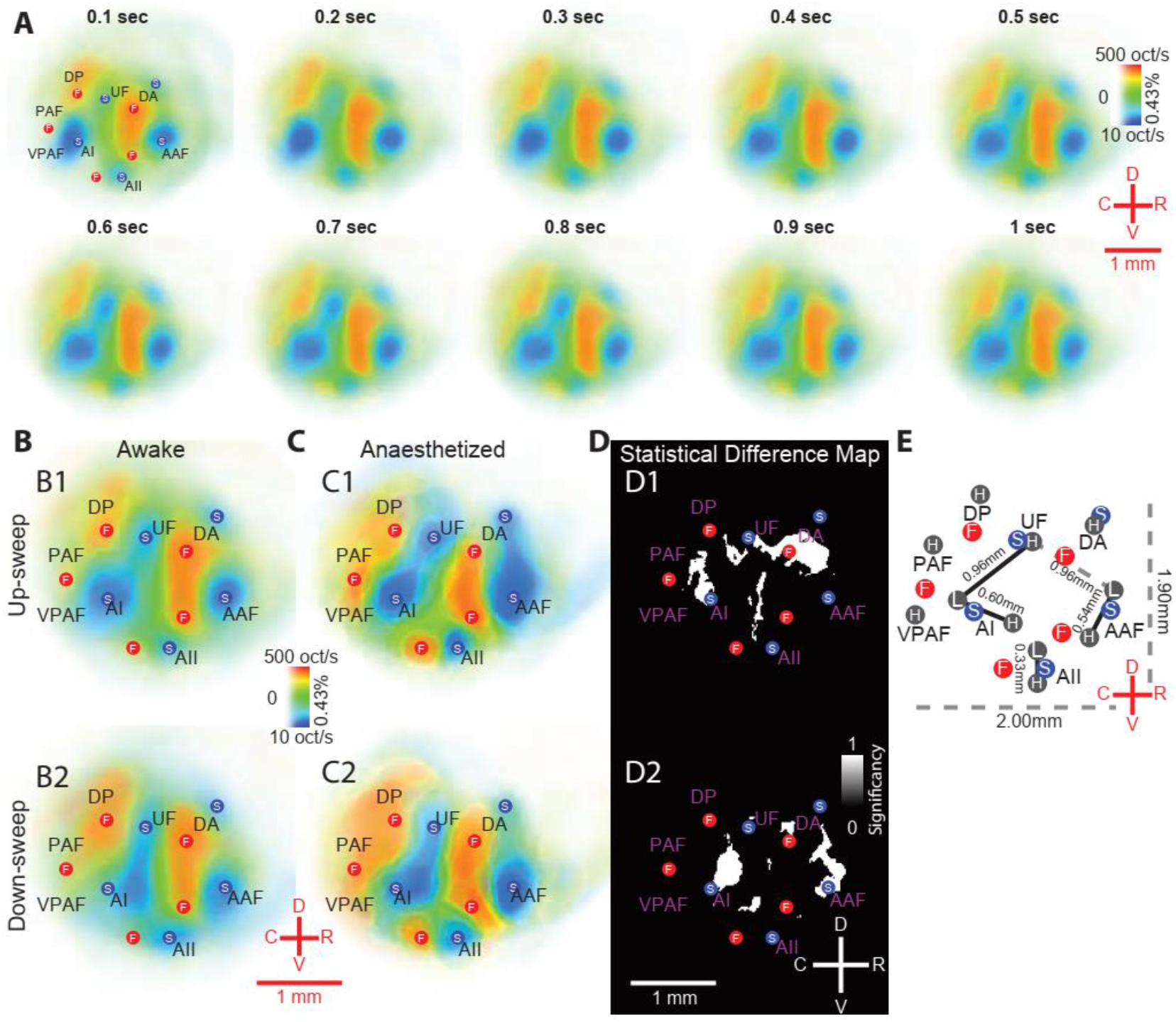
The auditory cortex maps in response to frequency modulations (FMs). **(A)** The average of 36 awake and 6 anesthetized upward FM sweep rate maps, were merged using landmarks registration (n = 12 mice, 3-4 months of age). To register all the maps to a common framework, slow (S) and fast (F) frequency sweep landmarks were first identified for each map and then the average of landmarks was determined. Maps from multiple recording sessions were aligned and merged using the transformation of landmarks to the average of landmarks. For the colormaps, the best sweep rate is indicated by color and the responsiveness to the sound by saturation. Maps were visually similar after 100 ms, with minimal differences across different time windows. Five regions associated with slow sweeps (AI, AII, AAF, DA, and UF) and five regions associated with fast sweeps (PAF, AII, AAF, UF, and DP) were distinguished, including some previously identified regions (AI, AII, AAF, and UF) and some regions that were newly identified in this study (DA, DP, and PAF). These slow and fast landmarks appear in both upward and downward sweep directions. For consistency and ease of comparison, only the landmarks of upward sweep frequency are shown. **(B** and **C)** Upward (B1 and C1) and downward (B2 and C2) FM sweep rate average maps for awake **(B)** and isoflurane-anesthetized **(C)** states in mouse auditory cortex, with a 0.5s averaging window beginning from stimulus onset. In downward compared to upward sweep rate maps, the center of the slow sweep rate region in AI is shifted 350 μm rostrally. **(D)** Maps of statistical analyses (using the Kruskal-Wallis test) comparing the best sweep rate between awake and anesthetized states for upward (D1) and downward (D2) sweeps. The white color shows the regions where the best sweep rate was different between the awake and anesthetized maps (p < 0.05). Major differences were observed at the two sites. For one site, at AI, the anesthetized maps showed lower values (i.e., darker blue color) of the best sweep rate compared to the awake maps. The second site was between AAF, DA, and UF, where anesthetized maps demonstrated the expansion of slower sweep rate areas and the shrinkage of faster sweep rate regions compared to the awake maps. All auditory areas cover less than 2×2 mm^2^. In general, the direction of the sweep rate gradient was from rostral to caudal. (E) The outline of tonotopy and FM sweep rate landmarks in the mouse auditory cortex averaged from a total of 42 awake and anesthetized recordings.

Similar to the linear regression analysis for pure tones, a linear regression model was fit to the FM stimuli, behavioral measures, and behavioral video as predictive features, and cortical activity as regressor output. The results of the statistical test of larger explained variance of FM stimuli compared to the explained variance of behavioral measures and video is shown in Figure S3A. Both maps of significant regions at p-value < 0.05 level (left) and p-value at each pixel (right) are displayed. The map of pixels that their R^2^ values for FM stimuli are greater than that of behavioral measures and video spans to include all the identified slow and fast sweep rate landmarks. Thus, this analysis emphasizes the high responsivity of these regions to FM stimuli.

Figure 4A demonstrates the reliability of the FM sweep rate maps at multiple time averaging windows after the response onset, ranging from 0.1-1 s. The maps have similar slow and fast rate landmark locations and FM sweep rate gradients. This finding shows that the auditory cortical responses to FM sweep chirps involve slow and stable dynamics after the response onset, similar to our previous finding for the cortical representation of pure tones over time. Figures 4B and 4C display the average FM sweep rate maps of awake and isoflurane-anesthetized states, respectively, after registration to a common coordinate framework. The top maps (B1 and C1) represent the upward sweep rate topographic maps, and the bottom maps (B2 and C2) show the downward sweep rate topographic maps. Despite similar layouts of slow and fast sweep landmarks and gradients in these four topographic maps, the maps also showed some differences. The locations of all landmarks in the upward sweep rate map were identical to the locations of the landmarks in the downward sweep rate map, except for the slow rate landmark in AI. The location of this slow rate landmark in the downward sweep rate map was approximately 350 μm rostral compared to its location in the upward sweep rate map. This difference between upward and downward sweep rate maps was found in both awake and anesthetized states. Differences between maps in awake and anesthetized states were also visually apparent in areas such as AI. To statistically compare the topographic maps between awake and anesthetized states, the Kruskal-Wallis test was applied to the distribution of best FM sweep rate maps in the two brain states for each of the upward and downward sweeps. The results of the statistical comparisons between awake and anesthetized FM sweep rate maps are plotted in Figure 4D, in which panels D1 and D2 show the statistical difference map for the upward and downward sweeps, respectively. The p-value maps corresponding to Figures 4D1 and 4D2 are plotted in Figures S3B and S3C. In Figure 4D, the white color represents the pixels for which the best FM sweep rate differed significantly between the two brain states, including one region located close to AI and another region spanning AAF, DA, and UF (median p-values in table S3D; AI: up-sweep p = 0.0472, down-sweep p = 0.0163; AAF-DA-UF: up-sweep p = 0.0163, down-sweep p = 0.0283). For the region near AI, the darker blue color in the anesthetized state map compared to the awake state map indicates that the best sweep rate was lower in the anesthetized state. In the second region, there were areas between AAF and DA in which higher best sweep rates (yellow and orange colors) in the awake state changed to lower best sweep rates (blue color) in the anesthetized state. In summary, despite the overall similarity between the sweep rate maps in the two states, slower sweep rates were more encoded in the anesthetized maps. Figure 4F demonstrates a schematic of the mouse auditory cortical regions and landmarks for both tonotopic and FM sweep rate maps obtained in this study.

### 3.4. The role of fast FM sweep rate regions in the cortical representation of mouse ultrasonic vocalization

Mice use USVs primarily in their social interactions to encourage or hinder social behaviors (Portfors, 2007). USVs are classified based on their time-frequency spectrogram shapes. In this study, 10 adult mouse USVs were used, each representing a distinct shape of the time-frequency spectrogram (Figure 5A) (Grimsley et al., 2011). The occurrence percentage of these vocalizations is age-dependent, and syllabi numbers 6 (upward frequency-modulated spectrum), 1 (flat spectrum), 4 (chevron), and 5 (downward frequency-modulated spectrum) are more frequent in adult mice. These representative USVs are rich in shape complexity, frequency range, and time duration, and often have a time duration between 5-50 ms, high-frequency spectrum, and fast sweep rates (Grimsley et al., 2011).

**Figure 5.**
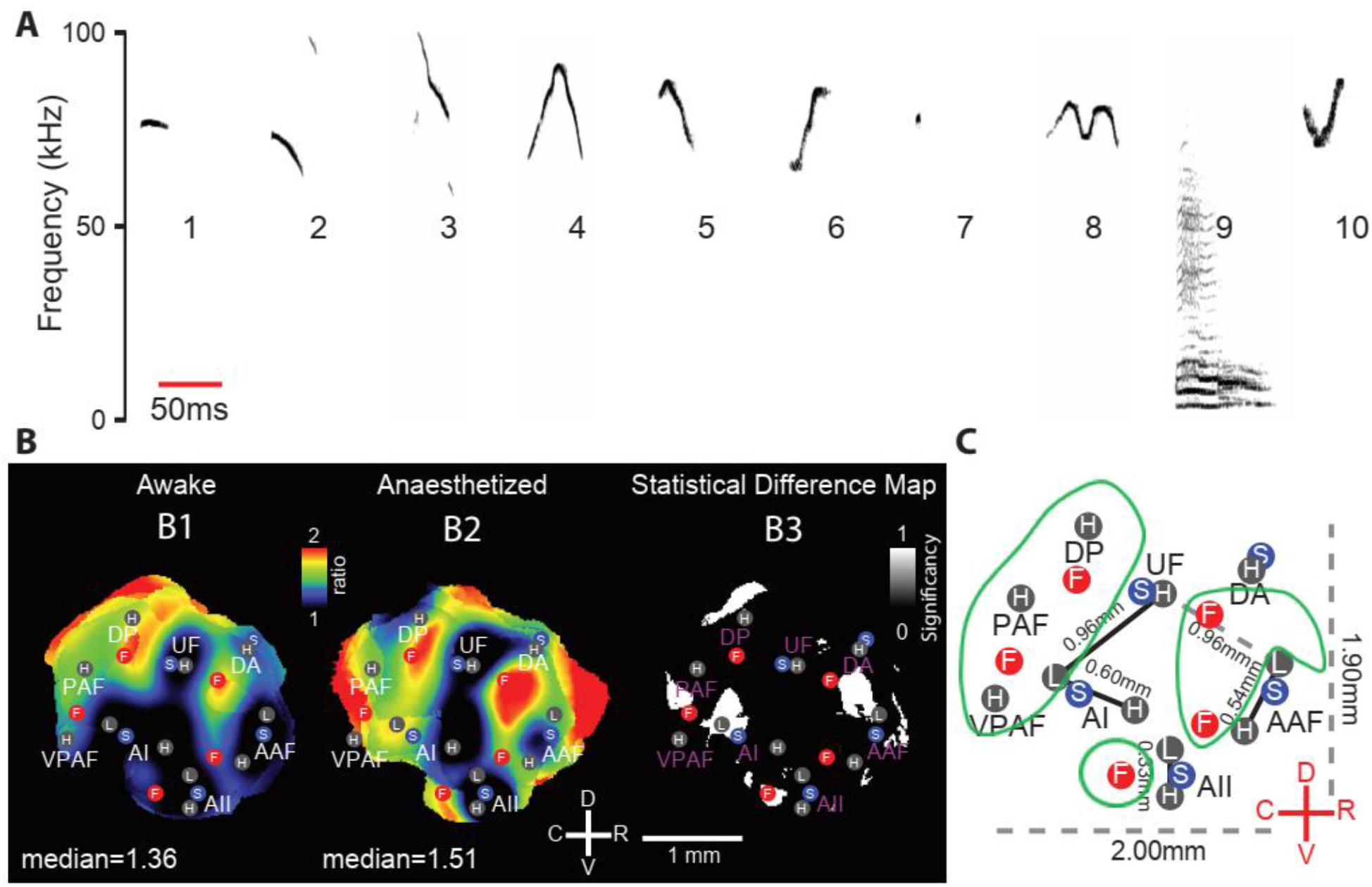
The response ratio of mouse ultrasonic vocalizations (USVs) to tonal stimuli (V/T ratio) in awake and isoflurane-anesthetized states. **(A)** The spectrogram of mouse USVs used in this study (Grimsley et al., 2011). The horizontal red bar shows the time scale of 50 ms. Each number from 1 to 10 corresponds to one type of USV. Each USV type was presented 30 times at 80 dB SPL level in random order. The responses were collected and then averaged over all days of recordings (3 days for awake and 1 day for anesthetized states) to get one single average response. The average response of auditory cortical regions to all tonal stimuli (4, 11.3, and 32 kHz) at 80 dB SPL level across all days of recordings (3 days for awake and one day for anesthetized states) was also determined. The V/T ratio was calculated for each animal. Then, using landmarks, all ratios were transformed into a common framework to get the average ratio map for all animals. **(B)** The V/T ratio in awake (B1) and isoflurane-anesthetized (B2) states for the auditory cortex window. In both states, vocal stimuli evoked stronger responses in areas corresponding with fast FM sweep rates; this finding was not observed in areas related to high frequencies. This finding means that those regions of the auditory cortex that process fast FM sweep rates are more responsive to vocal (complex) than tonal (simple) stimuli. **(C)** Map of statistical analyses (using the Kruskal-Wallis test) to compare the V/T ratio between awake and anesthetized states (n = 6). The V/T ratio was higher (p < 0.05) in four sites anesthetized compared to awake maps (AI, AII, AAF, and PAF). In DP the awake map showed higher values than the anesthetized map. **(D)** The outline of the auditory cortical regions with tonotopic (black circles) and FM sweep rate (blue and red circles) hotspots, as well as areas with a V/T ratio higher than one (green enclosures).

Before finding cortical regions that are more responsive to mouse USVs compared to pure tones, a linear regression analysis was performed to find regions that are more sensitive to USVs compared to behavioral measures and video. The contribution of USV stimuli in the construction of cortical activity was calculated and compared to the contribution of the behavioral measures and video. Figure S4A displays the p-value map (right) and the significant regions at the p-value < 0.05 level (left) in which the explained variance of USV stimuli is greater than that of behavioral measures and video. The map of significant regions shows that all identified regions and landmarks in the tonotopic and FM rate topographic maps are more sensitive to USVs compared to behavioral measures and video.

Next, to compare the cortical representation of vocal stimuli to pure tones, the ratio of the average response to all USVs divided by the average response to all pure tones (V/T) was computed at 80 dB SPL per animal and then averaged over all animals. Figure 5B shows these average V/T ratios for awake (B1) and isoflurane anesthetized (B2) states. The color bar reflects the value of the V/T ratio, in which hot colors refer to higher values and black color represents values less than or equal to 1. For reference, the locations of the AC tonotopic and FM upward sweep rate landmarks are overlaid on the V/T ratio maps. In both awake and anesthetized states, similar regions with a V/T ratio higher than 1 were distributed across the AC, including AII, DP, PAF, VPAF, and a region between UF and AAF. Most of these high V/T ratio regions were associated with the processing of fast FM sweep rates. The V/T ratios at two of these regions, the two fast sweep rate landmarks closest to the UF region, were the highest in both awake and anesthetized states. The median values of the V/T ratio for the regions favoring USVs (i.e., pixels with V/T > 1) were 1.36 and 1.51 for awake and anesthetized states, respectively. Conversely, for almost all the slow FM sweep rate landmarks the V/T ratio was less than or equal to 1 (note the black color at these landmarks). These regions include parts of AI, AII, AAF, and UF. Interestingly, despite the high-frequency content of USVs, high-frequency tonotopic regions did not contribute to the representation of USVs more than pure tones. The large black regions (V/T ≤ 1) in the middle of the maps included high-frequency regions of AI, AII, and UF. The V/T ratio also was affected by brain state; the extent of the regions responsive to USVs was greater in the anesthetized compared to the awake state. To quantify this difference, the Kruskal-Wallis test was carried out on the distribution of V/T ratios in awake and anesthetized states. Five regions (white color areas in Figure 5B3: AI, AII, AAF, DP, and PAF) showed significantly different (p-value < 0.05) V/T ratios between awake and anesthetized states (Figure 5B3). Figure S4B represents the corresponding p-value map, and table S4C exhibits the median p-values for the five regions with significant differences between awake and anesthetized states (median p-values: AI p = 0.0176, AII p = 0.0296, AAF p = 0.0285, DP p = 0.0330, and PAF p = 0.0339). Except for DP, the V/T ratio for the anesthetized state was larger than the awake recordings in these regions. This finding demonstrates the greater response of the AC to USVs than pure tones in the anesthetized compared to the awake state. Finally, Figure 5C illustrates the outline of the auditory cortical regions, with the main landmarks from the tonotopic and FM sweep rate maps and three green contours indicating regions with V/T ratios larger than 1.

### 3.5. The best frequency and FM sweep rate pattern are sigmoid functions of distance in auditory cortical maps

In this final component of the study, the relationship between best frequency and distance from the center of a low-frequency landmark in a tonotopic gradient was quantified. Similarly, the best FM sweep rate in a topographic gradient was characterized as a function of distance from the center of a slow FM sweep rate landmark. These characterizations provide metrics to estimate the best frequency or FM sweep rate of the desired location given the best frequency or FM sweep rate in other locations in the mouse AC. For each of the tonotopic gradients shown in Figure 6A, a line from the center of the low-frequency landmark to the center of the high-frequency landmark was drawn. Along this line, the best frequency and its corresponding distance from the low-frequency landmark were quantified for each animal. Figures 6B and 6C represent the average of the best frequency values (solid lines) and the standard error of the mean (shaded error bars) for each color-coded tonotopic gradient as a function of distance for both awake and isoflurane-anesthetized states. Each one of the tonotopic gradients showed a similar frequency span and trend of the best frequency as a function of distance in both states. Almost all graphs showed a similar shape resembling a sigmoid function [S-shape: frequency = 1/(1+e^distance^)]. Figures 6D and 6E show the inverse sigmoid function based on the best frequency values in each tonotopic gradient in awake and anesthetized states, respectively. The associations were highly linear (R^2^ > 0.93 and 0.90 for awake and anesthetized states, respectively) and confirmed the sigmoid relationship between the best frequency and distance from the low-frequency landmark in a tonotopic gradient. In the sigmoid functions, the higher and lower ends displayed a smoother frequency change compared to the middle frequencies as a function of distance. This finding suggests that low and high frequencies occupy longer distances compared to middle frequencies in a tonotopic gradient.

**Figure 6.**
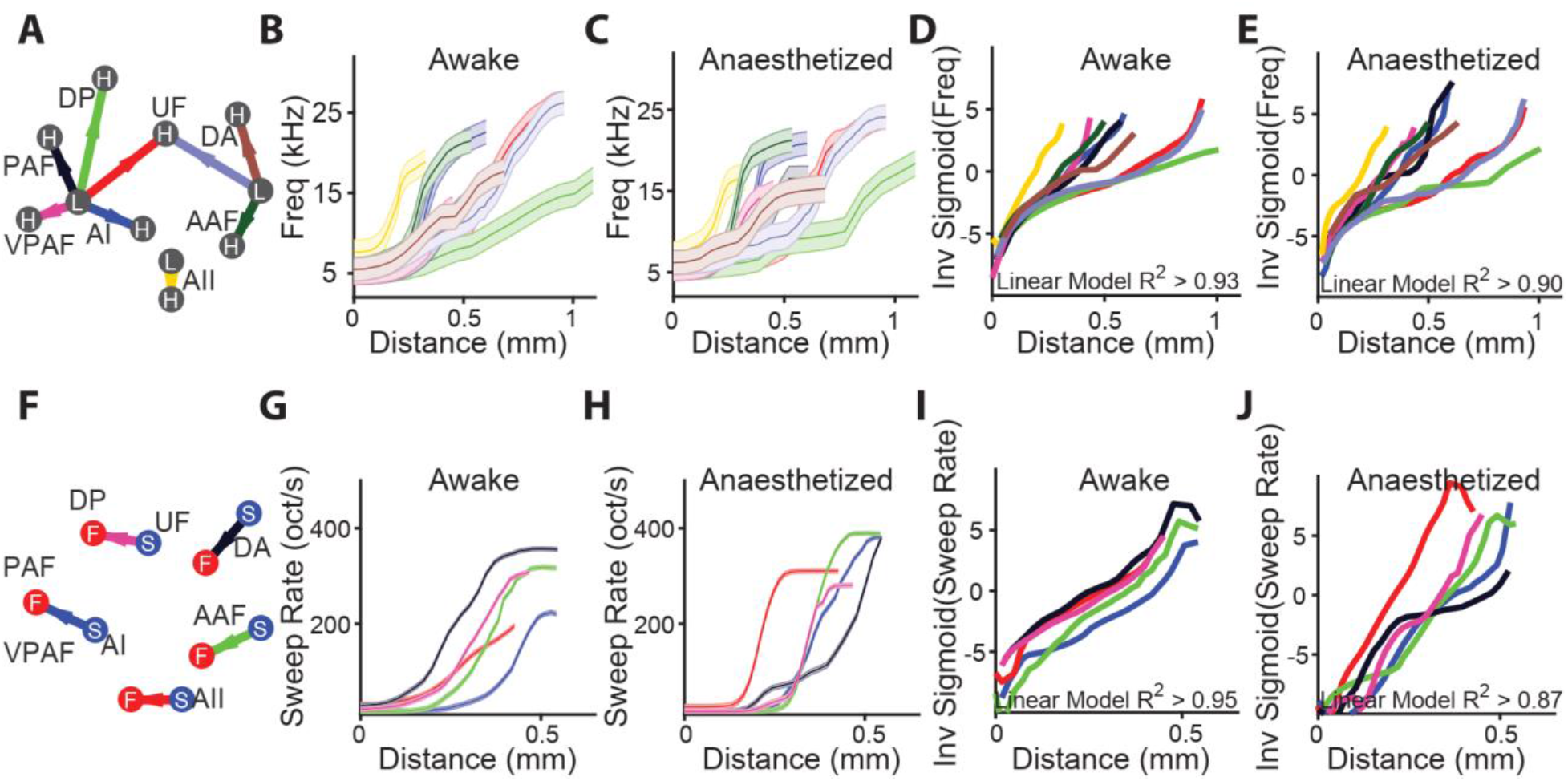
The alteration of the best frequency and the best FM sweep rate as a function of distance in tonotopic and FM sweep rate gradients of the mouse auditory cortex. **(A)** The frequency-distance association in tonotopic regions in awake (n = 12, 36 recordings) and isoflurane-anesthetized (n = 6, 6 recordings) mice. The graph shows the direction of tonotopic gradients from low to high-frequency landmarks in distinct colors. **(B** and **C)** The association between the best frequency and distance from the center of low-frequency landmarks in awake **(B)** and isoflurane-anesthetized **(C)** states. Each color corresponds to a tonotopic gradient shown in **A**. Solid lines exhibit the mean values for all animals, and the shaded areas reflect the standard error of the mean (SEM). The graphs resemble a sigmoid (S-shaped) function [frequency = 1/(1 + e^distance^)]. **(D** and **E)** The inverse-sigmoid graphs of the best frequency-distance association from low to high-frequency landmarks matched **B** and **C** respectively. The minimum adjusted R-squared values of a linear model fitted to the inverse-sigmoid graphs (0.93 and 0.90 for awake and anesthetized, respectively) demonstrate that the relation between the best frequency and distance was highly similar to a sigmoid function. The frequency alterations were steeper for middle frequencies compared to lower and higher frequencies, which indicates that larger distances in the tonotopic maps are occupied by low and high frequencies compared to middle frequencies. **(F-J)** Same as **A-E** but for the FM sweep rate gradients.

A similar relationship was observed between the best FM sweep rate and distance from the center of a slow landmark in an FM sweep rate gradient. Figure 6F shows the gradient lines from each slow landmark to its corresponding fast landmark. After calculating the best FM sweep rate along these gradients for each animal, the best sweep rate values were plotted relative to the distance in Figures 6G and 6H for awake and anesthetized states, respectively. A sigmoid function was obtained in the plots (Figures 6I and 6J for awake and anesthetized, respectively) after fitting an inverse-sigmoid function to each graph (R^2^ > 0.95 and 0.87 for awake and anesthetized states, respectively). These results point to a common underlying property of cortical encoding of sound stimuli features, in which representation of changes in a feature is non-uniformly distributed over the auditory cortical surface.

## 4. Discussion

This study aimed to precisely distinguish the auditory cortex topographic maps in response to simple and complex auditory stimuli in both awake and isoflurane-anesthetized states. The six major findings were: 1) Three newly identified tonotopic regions were generally responsive to high-frequency tonal stimuli. 2) The tonotopic maps showed high reproducibility over time within and between animals. 3) In the awake versus anesthetized state, the amplitude of tonotopic maps was greater in response to high frequencies (i.e., better hearing sensitivity). 4) In the anesthetized state, the response map to faster FM sweep rates was shrunk relative to the awake state. 5) A stronger response to USVs compared to tonal stimuli was observed in areas associated with faster FM sweep rates. 6) Modeling the frequency-distance association in tonotopic gradients and sweep rate-distance association in FM sweep rate gradients revealed a sigmoid function. In the following, we will consider these findings in turn.

### 4.1. Increased responsivity to higher frequencies in tonotopic maps

In this study, in addition to already known auditory cortical areas (i.e., AI, AII, AAF, DP, UF) (Guo et al., 2012; Issa et al., 2014; Tsukano et al., 2015), three new tonotopic regions (i.e., DA, PAF, VPAF) were characterized. The newly identified regions were generally responsive to higher frequencies and were in dorsal and caudal AC regions. This finding suggests that the AC tonotopic maps are oriented more toward high frequencies than low frequencies, which is consistent with the wider spectrum of high frequencies in the mouse hearing range (Heffner, 1980). The overrepresentation of behaviorally relevant frequencies (i.e., higher frequencies) in central auditory structures, including the AC, has previously been shown in several species including mice (Garcia-Lazaro et al., 2015). The tonotopic maps also were highly reproducible in the present study, which was examined in three different ways, including during a recording session, across three consecutive weeks of recordings, and between animals. Except for a few sparse pixels, the tonotopic maps were quite similar over time and between animals. The strong reproducibility of tonotopic maps suggests that potential confounding factors, such as environmental or technical issues, had little impact on the results.

### 4.2. Better hearing sensitivity at high frequencies in awake vs. anesthetized maps

In this study, the response amplitude at high frequencies (the H region of AI and UF) was greater in the tonotopic maps from awake compared to anesthetized mice. This finding implies that hearing sensitivity at high frequencies is diminished in the anesthetized relative to the awake state. In this study, isoflurane (1.0-1.5%) anesthesia was used. It has been shown that higher dosages of isoflurane, greater than 1.6%, can significantly lower auditory thresholds (Sheppard et al., 2018). Isoflurane is a fast-acting anesthetic that is widely used in experimental research. Several studies demonstrate the impact of isoflurane on both peripheral (Stronks et al., 2010; Cederholm et al., 2012; Sheppard et al., 2018) and central (Stronks et al., 2010; Ruebhausen et al., 2012; Bielefeld, 2014) auditory systems. For instance, isoflurane can decrease cochlear amplification (Stronks et al., 2010) and the amplitude of otoacoustic emissions (OAE) (Sheppard et al., 2018) and auditory brainstem responses (ABR) (Stronks et al., 2010). The possible mechanisms underlying these impacts are changes in cochlear blood flow (CBF) (Nakashima et al., 2003; Xiao et al., 2014), the blockage of acetylcholine (ACh) receptors (Dilger et al., 1992), and enhanced gamma-aminobutyric acid (GABA)-gated channel currents (Ming et al., 2001), which lead to increased inhibition mediated by medial olivocochlear (MOC) efferents (Sheppard et al., 2018). In the present study, a low dosage of isoflurane was used to model the impact of sleep state on mouse tonotopic maps. Few studies have examined the effect of isoflurane and/or natural sleep on mouse tonotopic maps’ response amplitudes. Replication of our findings during natural sleep will clarify whether changes in hearing sensitivity result from using isoflurane or are associated with sleep processes.

Natural sleep is considered a powerful model to study the neuronal correlates of changes in brain state (Wilf et al., 2016). There is a mutual association between sensory systems and the central nervous system supporting the awake state. It was found that sensory systems show electrophysiological changes that are associated with brain state (Velluti, 1997). Both animal electrophysiological studies (Steriade et al., 1991; Coenen and Drinkenburg, 2002) and human evoked/event-related potential (AEP/ERP) findings (Hennevin et al., 2007; Wilf et al., 2016) indicate little state-dependent change in prethalamic transmission of auditory signals (Portas et al., 2000). But at the thalamocortical level, significant sleep-related modulation of the latency and/or amplitude of auditory responses or single neurons’ firing rates have been reported (Bastuji et al., 2002; Coenen and Drinkenburg, 2002; Issa and Wang, 2008, 2011). For example, no changes were shown in human auditory early-latency responses (originating from the acoustic nerve and brainstem) during non-rapid eye movement (NREM) sleep (Amadeo and Shagass, 1973; Mendel and Kupperman, 1974; Bastuji et al., 1988), altered auditory middle-latency responses (originating from the auditory thalamus and primary auditory cortex) (Erwin and Buchwald, 1986; Deiber et al., 1989), as well as auditory late latency responses (cortical in origin), were reported (Wesensten and Badia, 1988; Brualla et al., 1998; Cote and Campbell, 1999; Perrin et al., 1999). Electrophysiological studies in cats also show a decline in neuronal firing rates in the parietal association cortex (Steriade et al., 1978) and the thalamus (Steriade and Hobson, 1976) during NREM sleep. In conclusion, these findings support changes in thalamocortical auditory processing in response to reduced conscious perception during sleep (Portas et al., 2000). The impact of sleep compared to anesthetized and awake states should be studied in the future.

### 4.3. Differences between FM sweep rate maps in anesthetized vs. awake state and upward vs. downward sweep directions

In this study, for both upward and downward sweep rates, five auditory cortical regions related to slow sweep rates (i.e., AI, AII, AAF, DA, and UF) and five related to fast sweeps (i.e., PAF, AII, AAF, UF, and DP) were distinguished, including four previously reported areas (i.e., AI, AII, AAF, and UF) (Stiebler et al., 1997; Honma et al., 2013; Tsukano et al., 2013; Tsukano et al., 2015) and three regions newly identified in our study (i.e., DA, DP, and PAF). The awake and anesthetized maps were statistically different in two tonotopic regions. The first difference was observed in AI, in which the best sweep rate was reduced in the anesthetized map compared to the awake state. The second difference was shown in the AAF-DA-UF regions, in which the response to slower sweep rates was expanded and the response to faster sweep rates was reduced in maps from the anesthetized compared to the awake state. AAF is specialized for faster temporal processing relative to AI and DA (Trujillo et al., 2011), and the shrinkage of a faster sweep rate map in this region points to the reduced processing of stimuli associated with faster sweep rates, such as vocal stimuli, in the anesthetized versus the awake state.

In this study, a difference between upward and downward FM sweep maps also was observed. Despite the report of no difference between upward and downward FM sweep maps in a previous study (Issa et al., 2017), we found that the center of the slow sweep rate responsive area in AI shifts 350 μm rostrally between upward and downward FM sweep maps. This finding suggests that the best FM sweep rate of associated neuronal populations differs for upward and downward sweeps. Sweep direction determination contributes to the detection of complex sounds such as USVs (Razak and Fuzessery, 2006; Aponte et al., 2021). This region might be analogous to a previously identified region in rat AC where a topographic map of FM sweep direction was found (Zhang et al., 2003). Future research is necessary to reveal the function of this region at the cellular level in encoding sweep direction for simple and complex sounds.

### 4.4. Higher V/T ratio in regions associated with the processing of faster FM sweep rates

Little is known about the AC regions that respond to USVs (50-100 kHz) in rodents (Carruthers et al., 2013; Issa et al., 2014). In our study, vocal stimuli elicited a stronger response than tonal stimuli in both anesthetized and awake states, and a higher V/T ratio was obtained in the regions (AII, DP, PAF, VPAF, and a region between UF and AAF) linked to the processing of faster FM sweep rates compared to regions that respond to high frequencies. This finding suggests that tonotopic regions related to faster sweep rates make a greater contribution to vocal processing than tonotopic regions related to higher frequencies. The V/T ratio was larger in four sites in anesthetized versus awake maps (AI, AII, PAF, and a region next to AAF), as well as in the DP region in awake compared to anesthetized maps.

In this study, a craniotomy was made only over the right AC. Previous findings demonstrate that mice process intra-species vocalizations preferentially in the left AC (Ehret, 1987; Marlin et al., 2015), and the right AC contributes to more general auditory computational tasks, such as the detection of sweep direction (Zhang et al., 2003; Wetzel et al., 2008). For instance, in a recent study using 3D volume imaging of fos expression, sweep stimuli led to extensive activation of the right AC, whereas USVs preferentially activated the anterior regions of the left AC, where ultrasonic frequencies are represented (Levy et al., 2019). Future studies using craniotomies over left and right AC and dorsal cortical areas can shed more light on the lateralization of vocal processing in the auditory and associative cortices and the processing of fast sweep rates in the mouse brain.

### 4.5. A sigmoid frequency-distance function in tonotopic maps

The mouse hearing frequency range extends from 2.3 to 92 kHz (at an intensity of 60 dB SPL), demonstrating a V-shaped audiogram (Heffner, 1980). Hearing sensitivity gradually increases and shows higher sensitivity (below 10 dB SPL) at middle frequencies between 8-32 kHz. The best hearing threshold appears near 16 kHz with an average threshold of −10 dB SPL. Above 16 kHz, the hearing sensitivity begins to drop gradually with a steeper decline above 64 kHz (Ehret, 1977; Heffner, 1980). In this study, our investigation of the frequency-distance association in tonotopic maps showed a sigmoid function in both awake and anesthetized states. This sigmoid function reflects steeper frequency changes in a shorter distance for middle frequencies versus lower and higher frequencies. Our findings support the idea that the middle frequency area of AC tonotopic maps, corresponding with the area of best hearing sensitivity and frequency discrimination in mice (Ehret, 1977; de Hoz and Nelken, 2014), is smaller but with higher frequency density and faster frequency variation compared to tonotopic areas before (responding to lower frequencies) and after (responding to higher frequencies) this region. In summary, our model of sigmoid frequency-distance functions in tonotopic maps is consistent with behavioral evidence of a V-shaped audiogram and better hearing sensitivity at middle frequencies (i.e., 8-32 kHz) in mice.

## 5. Conclusions

In this study, using wide-field Ca^2+^ imaging in Thy1-GCaMP6s mice, new topographic gradients for both tonal stimuli and FM sweep chirps were distinguished, in addition to previously identified AC maps. Our findings indicate the impact of both brain states (i.e., awake vs. anesthetized) and the complexity of auditory stimuli on mouse AC tonotopic maps. For instance, better hearing sensitivity at high frequencies and a stronger response to the best FM sweep rate were found in the awake compared to the anesthetized state. The response ratio of vocal to tonal stimuli also was larger in topographic regions associated with faster sweep rates. All tonotopic gradients also displayed a frequency-distance relationship representing a sigmoid function. The function of newly identified tonotopic maps in vocal processing and animal behavior, as well as the impact of natural sleep versus awake and anesthetized states on the AC and dorsal cortical areas, are relevant areas of research that should be investigated in the future.

## Acknowledgments

The present article was supported by Canadian Institutes of Health Research (CIHR) Grant# 390930, Natural Sciences and Engineering Research Council of Canada (NSERC) Discovery Grant #40352, Alberta Innovates (CAIP Chair) Grant #43568, Alberta Alzheimer Research Program Grant # PAZ15010 and PAZ17010, and Alzheimer Society of Canada Grant# 43674 to MHM; and a Canadian Institute for Advanced Research Grant 33033 to BK. This study was part of a Ph.D. dissertation project in Neuroscience that was approved by the Canadian Center for Behavioural Neuroscience (CCBN) at the University of Lethbridge under NSERC CREATE in BIP doctoral fellowship to N.A. We thank Dr. Brendan McAllister for inputs on the manuscript, Dr. Daniel Polley and Dr. Edward Hight for sharing their head-plate design, and Behroo Mirza Agha, Megan Sholomiski, Breanne Beatty, and Di Shao for assistance with experiments and animal husbandry.

## Author Contribution

Conceptualization, N.A., M.H.M., Z.J., and M.K.; Methodology, N.A., M.H.M., Z.J., J.S., and M.K.; Formal Analysis, N.A.; Writing - Original Draft, N.A., and Z.J..; Writing - Review & Editing, M.H.M., N.A., and Z.J.; Funding Acquisition, Resources, and Supervision, M.H.M.

---

**Figure S1.**
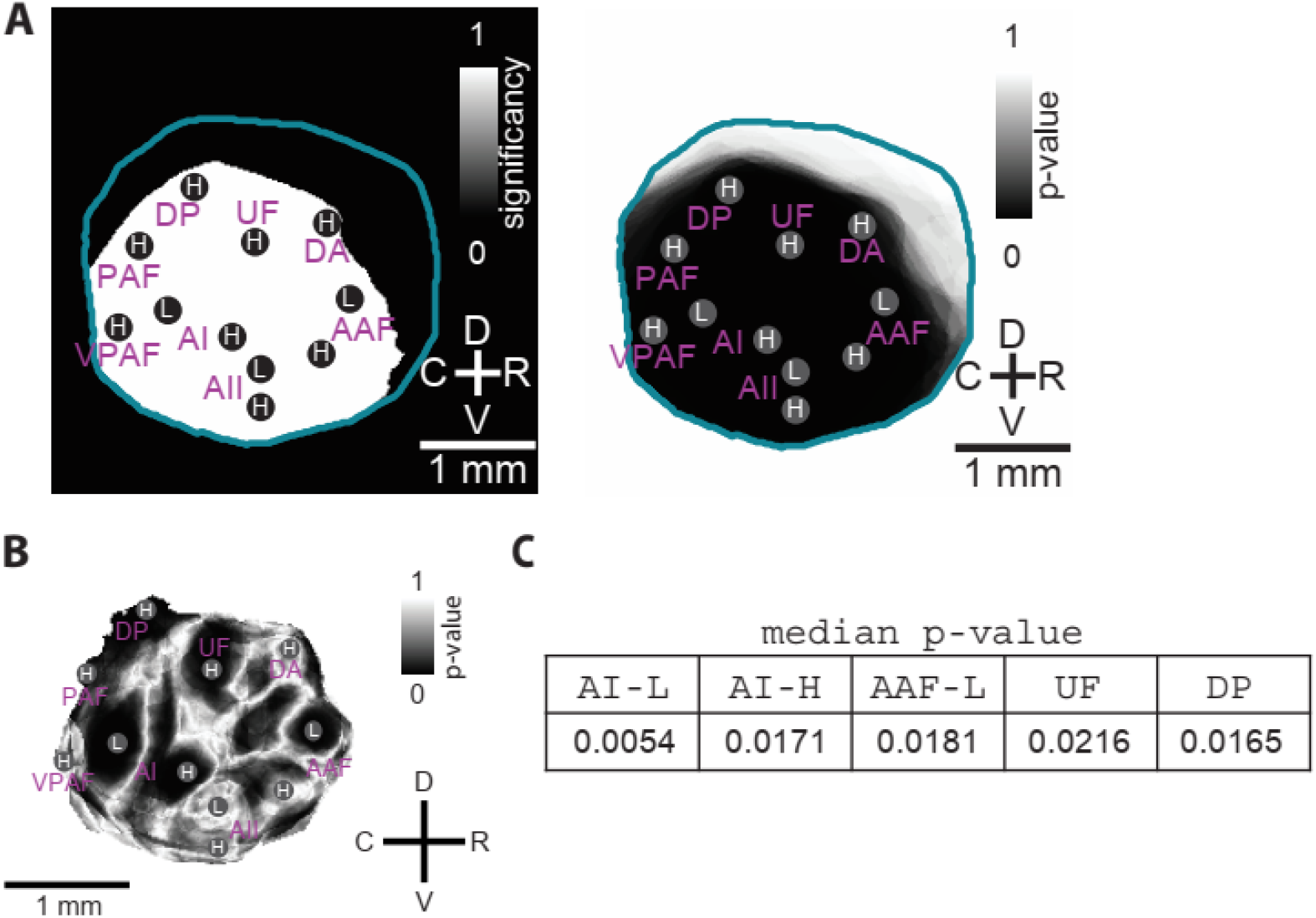
**(A)** The results of the statistical test (Wilcoxon rank-sum test at each pixel) of whether the explained variance of pure tone stimuli in the construction of wide-field cortical activity is greater than the explained variance of the behavioral measures and video. The left panel shows the significant regions at p-value < 0.05 and the right panel displays the corresponding p-value map. **(B)** The p-value map of statistical comparison (Kruskal-Wallis test at each pixel) on the best frequency (i.e., the highest response to L or H frequencies) between awake and anesthetized tonotopic maps. The corresponding significance map at the 0.05 level is plotted in Figure 2D. **(C)** Table of median p-values in five regions that show a significant difference in the best frequency between awake and anesthetized tonotopic maps. AI, primary auditory cortex; AAF, anterior auditory field; AII, secondary auditory field; DA, dorsal anterior field; DP, dorsal posterior field; PAF, posterior auditory field; UF, ultrasonic field; VPAF, ventral posterior auditory field.

**Figure S2.**
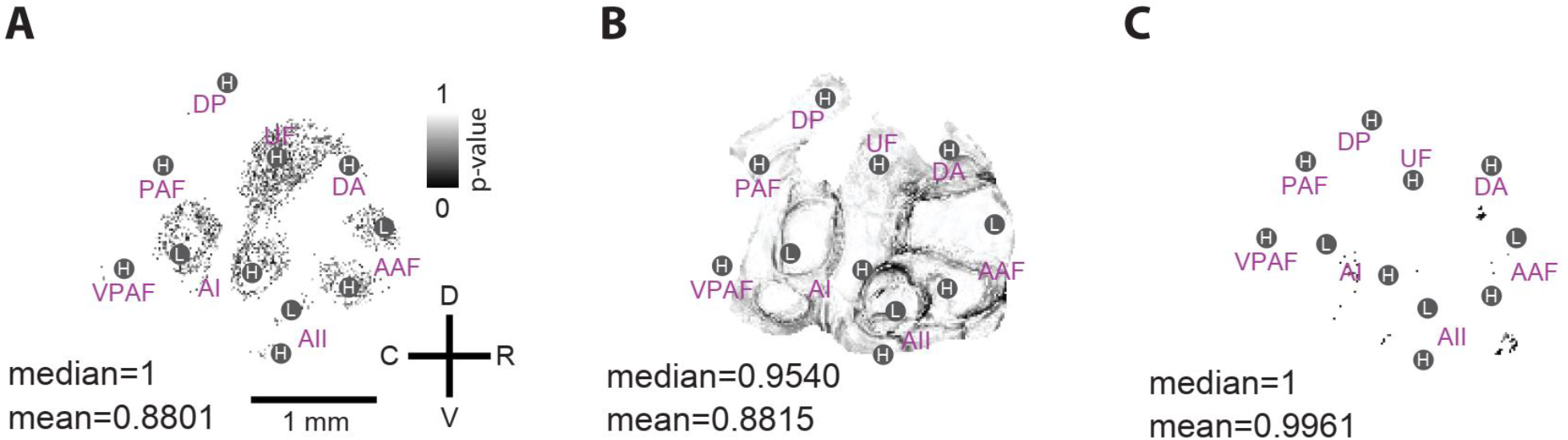
**(A)** The p-value map of the Chi-squared goodness-of-fit test on the best frequency distribution in a session of recording (corresponds to Figure 3A4). **(B)** The p-value map of statistical comparison (Kruskal-Wallis test) on the best frequency within the three weeks (corresponds to Figure 3B4). **(C)** The p-value map of the Chi-squared goodness-of-fit test on the best frequency distribution of all awake maps (corresponds to Figure 3C4). AI, primary auditory cortex; AAF, anterior auditory field; AII, secondary auditory field; DA, dorsal anterior field; DP, dorsal posterior field; PAF, posterior auditory field; UF, ultrasonic field; VPAF, ventral posterior auditory field.

**Figure S3.**
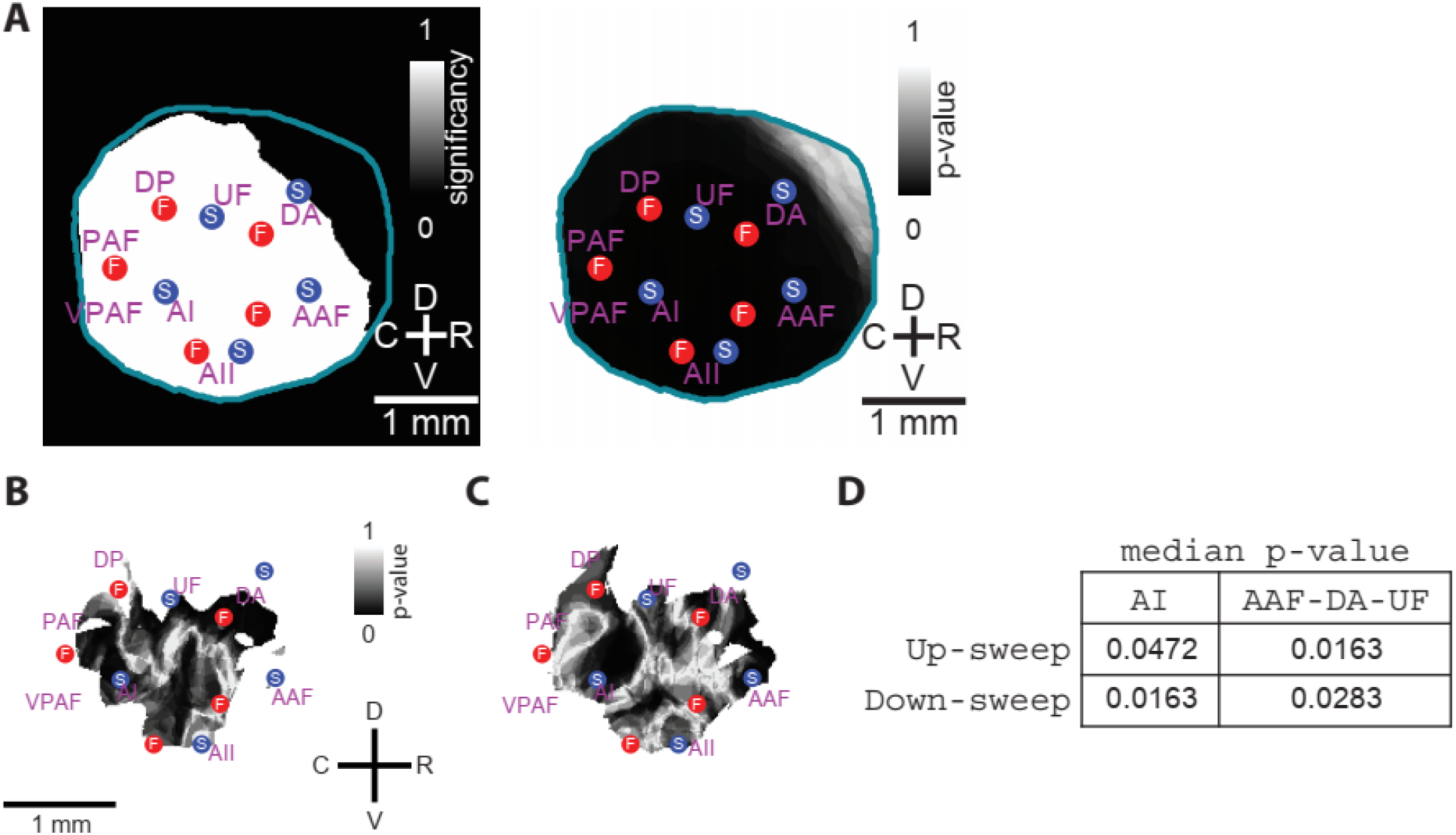
**(A)** The results of the statistical test (Wilcoxon rank-sum test at each pixel) of whether the explained variance of FM stimuli in the construction of wide-field cortical activity is greater than the explained variance of the behavioral measures and video. The left panel shows the significant regions at p-value < 0.05 and the right panel displays the corresponding p-value map. **(B** and **C)** The p-value maps of statistical analyses (using the Kruskal-Wallis test) on FM sweep rate between awake and anesthetized states for upward **(B)** and downward **(C)** sweeps correspond to Figures 4D1 and 4D2 respectively. **(D)** Table of the median p-values for the two regions that show a significant difference in the best FM sweep rate between awake and anesthetized states in both upward and downward sweep directions. AI, auditory cortex; AAF, anterior auditory field; AII, secondary auditory field; DA, dorsal anterior field; DP, dorsal posterior field; PAF, posterior auditory field; UF, ultrasonic field; VPAF, ventral posterior auditory field.

**Figure S4.**
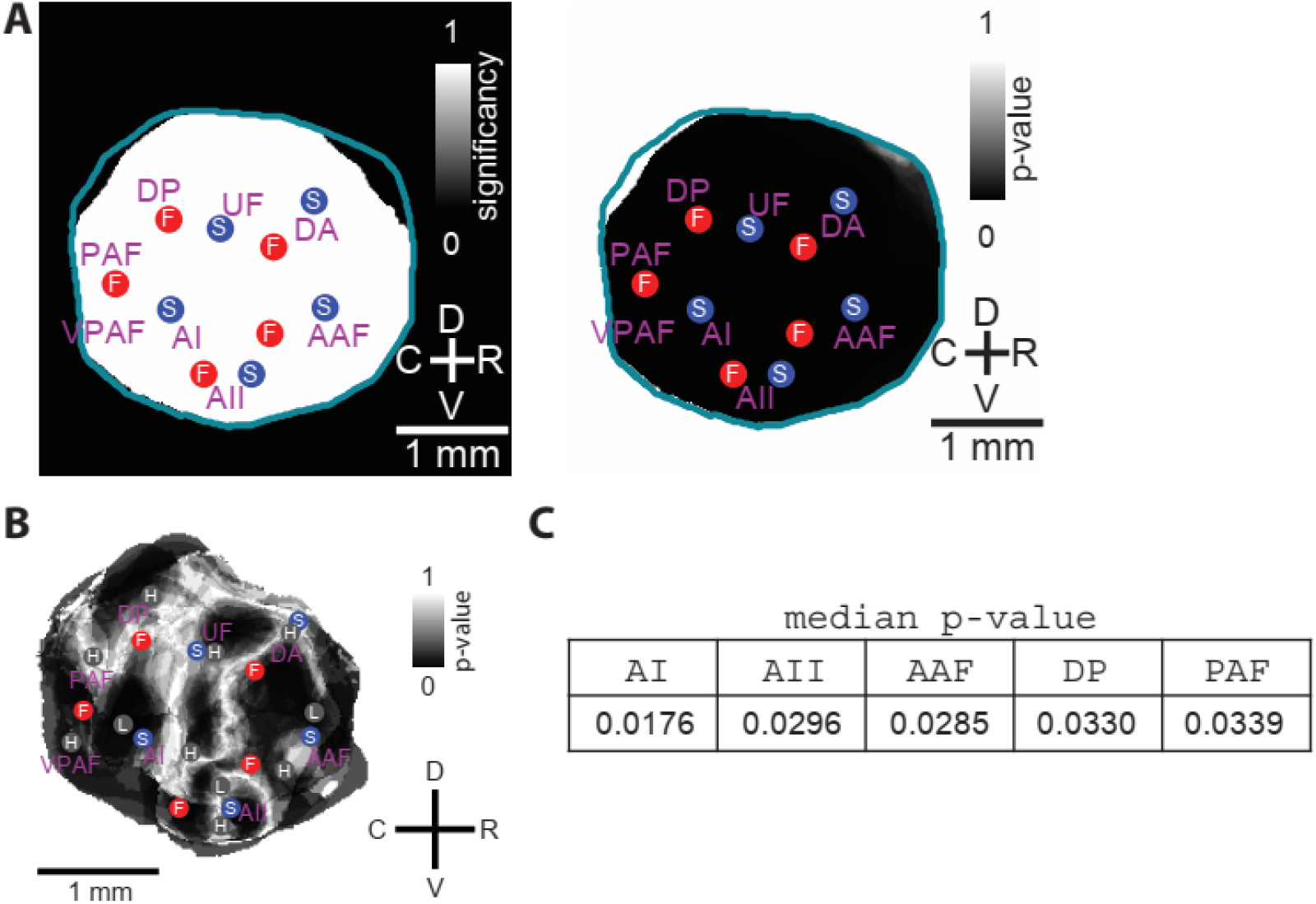
**(A)** The results of the statistical test (Wilcoxon rank-sum test at each pixel) of whether the explained variance of mouse USV stimuli in the construction of wide-field cortical activity is greater than the explained variance of the behavioral measures and video. The left panel shows the significant regions at p-value < 0.05 and the right panel displays the corresponding p-value map. **(B)** The p-value map of statistical analyses (using the Kruskal-Wallis test) to compare the V/T ratio between awake and anesthetized states (n = 6) (corresponds to Figure 5B3). **(C)** Table of median p-values for the five regions that show a statistically significant difference in V/T ratio between awake and anesthetized states. AI, auditory cortex; AAF, anterior auditory field; AII, secondary auditory field; DA, dorsal anterior field; DP, dorsal posterior field; PAF, posterior auditory field; UF, ultrasonic field; VPAF, ventral posterior auditory field.

